# Direct measurement of *in vitro* response to Praziquantel in *Schistosoma mansoni* populations from Western Kenya

**DOI:** 10.64898/2026.07.21.739336

**Authors:** Eric M. Ndombi, John Oguso, Peter Olilah, Churchill Orao, Brian Otieno, Madison Morales, Winka Le Clec’h, Frédéric D. Chevalier, Timothy Anderson

## Abstract

Large-scale treatment with praziquantel (PZQ) monotherapy is used to control schistosomiasis, leading to concerns about the emergence of PZQ-resistance. In Western Kenya, schistosome-infected patients frequently remain egg-positive following PZQ treatment, and several “hotspot” villages have been observed where transmission remains high, despite annual mass PZQ treatments. This project asks (i) whether PZQ-resistant parasites are found in Western Kenya and (ii) whether “hotspot” villages can be explained by a higher prevalence of PZQ-resistant parasites. We established a simple platform for directly assaying worm motility following *in vitro* PZQ-exposure in adult schistosomes isolated from a field setting. To do this, we established snail and hamster breeding colonies, and generated large populations of field-derived adult worms, by (i) harvesting *S. mansoni* eggs from multiple infected patients; (ii) infecting *Biomphalaria* spp snails with miracidia; (iii) infecting hamsters with released cercariae; (iv) perfusing adult worms from hamsters, and (iv) examining drug response following exposure to PZQ (1 µg/ml for 1 day) in individual *S. mansoni* worms using an automated movement assay. We measured PZQ-response in 1,800 adult male parasites, representing an estimated 185 parasite genotypes. We identified a single worm that remained motile after PZQ-exposure among the 185 parasite genotypes surveyed (frequency = 0.54%; 95% CI 0.01 - 2.97%, exact binomial) consistent with PZQ-resistant worms being extremely rare or absent. Our direct phenotypic screening results suggests that (i) PZQ-resistance is not currently an obstacle for *S. mansoni* control in Western Kenya, and (ii) that other factors explain the existence of persistent hotspots.

## INTRODUCTION

Mass drug administration using a single drug praziquantel (PZQ) has been the central strategy for the control of schistosomiasis since 1972. Almost 100 million people received PZQ treatment worldwide as preventive chemotherapy (Santé, 2025)with the aim of increasing this to 250 million doses per year. Mass treatment with PZQ has reduced schistosome transmission, and this reflected in lower prevalence of people with heavy infections and consequent reduction in clinical disease. However, the increasing coverage of PZQ monotherapy in targeted populations raises serious concerns about the evolution and spread of drug resistance in schistosome populations.

To date, there is no unambiguous evidence for PZQ resistance in natural schistosome populations, but there are multiple lines of evidence that make this a concern. First, several laboratory studies have demonstrated that resistant parasites can be selected in the laboratory. Parasites showing a (3-5-fold) change in drug response have been selected by PZQ treatment of either adult parasites in rodent hosts (Cioli et al., 2004; Fallon & Doenhoff, 1994; Pica-Mattoccia & Cioli, 2004) or larval parasites in the intermediate snail host (Couto et al., 2011). These experiments demonstrate that PZQ-resistance (PZQ-R) is a heritable genetic trait. In one case, the PZQ-resistance was mapped to a transient receptor potential channel (TRPM_PZQ_) that is now known to be critical for the mode of action of PZQ (Le Clec’h et al., 2021; Marchant, 2024). Furthermore, Le Clec’h et al (Le Clec’h et al., 2021) used marker assisted selection to purify populations of PZQ-R and PZQ-S parasites, which showed a 377-fold difference in EC_50_. Second, in both Egypt and Western Kenya - the location for this study - schistosome infected patients have been found that were not cured after multiple rounds of treatment (Ismail et al., 1999; Melman et al., 2009). Subsequent laboratory studies demonstrated that parasite isolates from these patients showed ∼5-fold increase in resistance to treatment demonstrating that treatment failures are due to parasite factors (Cioli et al., 2004). Third, regions where schistosome infected patients remain at high prevalence and intensity despite multiple rounds of PZQ-treatment have been observed in several countries (Assaré et al., 2020; Kittur et al., 2020). Such hotspot villages have been extensively studied in Western Kenya, the location of the current study (Kittur et al., 2020). A recent epidemiological study by our groups confirmed that these hotspots were maintained for eight years despite ongoing school-based deworming program (Olilah et al., 2026). The reasons for these “persistent hotspots” (Wiegand et al., 2017) are uncertain, but one possible explanation is that these represent foci of emerging PZQ resistance.

Two metrics – cure rate (CR) and egg reduction ratio (ERR) have been used to evaluate treatment efficacy of PZQ. However, two factors make these assays insensitive for detecting emerging PZQ-R in schistosome populations. First, populations of adult worms within hosts are genetically diverse, and only a fraction of parasite genotypes present may show PZQ-R. Second, PZQ is by no means a perfect drug: cure rates range from 60-90% in most endemic situations (King et al., 2011). A major deficiency is that immature schistosomes are refractory to PZQ treatment: in mice, 30 times more drug is required to kill 28-day old worms compared with 49-day old worms (Pica-Mattoccia & Cioli, 2004). Hence, it is difficult to determine whether treatment failure results from PZQ resistance or establishment of newly matured adult parasites (Danso-Appiah & De Vlas, 2002). This problem is further compounded, because assessment of cure rates relies on use of parasite egg counts, rather than direct measures of adult worm numbers before and after PZQ treatment. While there is a positive relationship between worm numbers and egg counts, the relationship is noisy (Cheever et al., 1977), and there are high levels of temporal variation in egg production when individual patients are sampled on consecutive days (De Vlas et al., 1992; Engels et al., 1997; Utzinger et al., 2001). Hence ERR that compare egg counts before and after PZQ treatment provide relatively insensitive metrics for detecting changes in PZQ treatment success and should only be able to detect PZQ resistance at high frequency within populations.

For protozoan parasites such as malaria, as well as fungal and bacterial pathogens, *in vitro* assays of drug efficacy can be used to empirically measure drug response in pathogen clones isolated from patients (Woodrow et al., 2013). This allows direct measurement of pathogen drug response phenotypes, without interference from host immunity, drug metabolism, and other factors that can cloud interpretation of therapeutic treatment studies. We aimed to develop a comparable method for direct *in vitro* measurement of frequencies of PZQ-R schistosome parasites from endemic areas. Here, we describe development of a platform for direct assessing frequencies of phenotypic PZQ-resistance in genetically diverse populations of *in vitro* cultured adult *Schistosoma mansoni* worms originating from infected patients in Western Kenya. We then examined PZQ-response in individual worms cultured in *in vitro* following exposure to PZQ using an automated movement assay.

## RESULTS

Fig. 1 provides an overview of our methods for generating large numbers of adult schistosome parasites for drug response assays.

**Figure 1.**
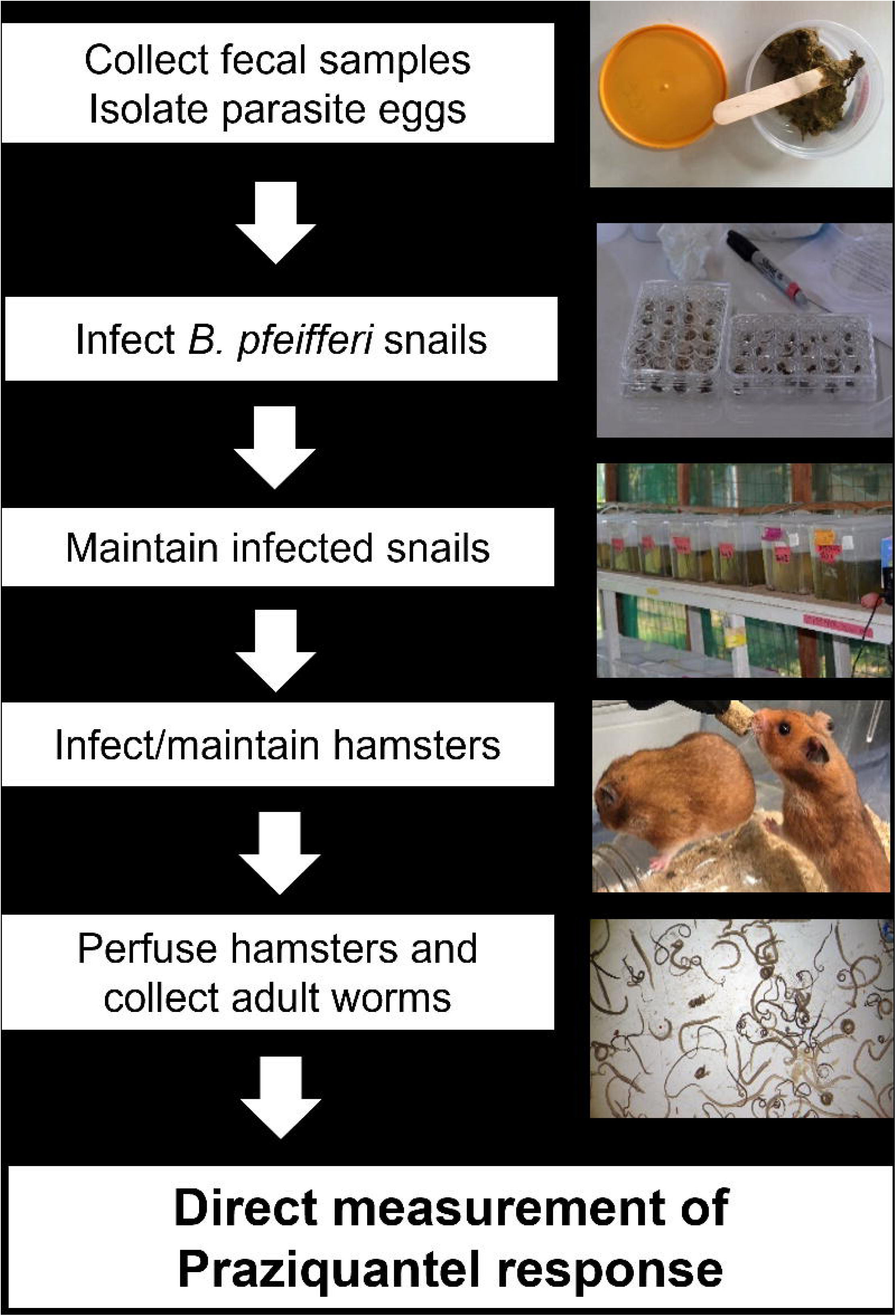
Workflow for producing genetically diverse field-derived adult parasites. Production of field derived adult parasites was conducted on site in Kisumu at the Kenya Medical Research Institute’s Centre for global Health Research, Kenya.

**Figure 2.**
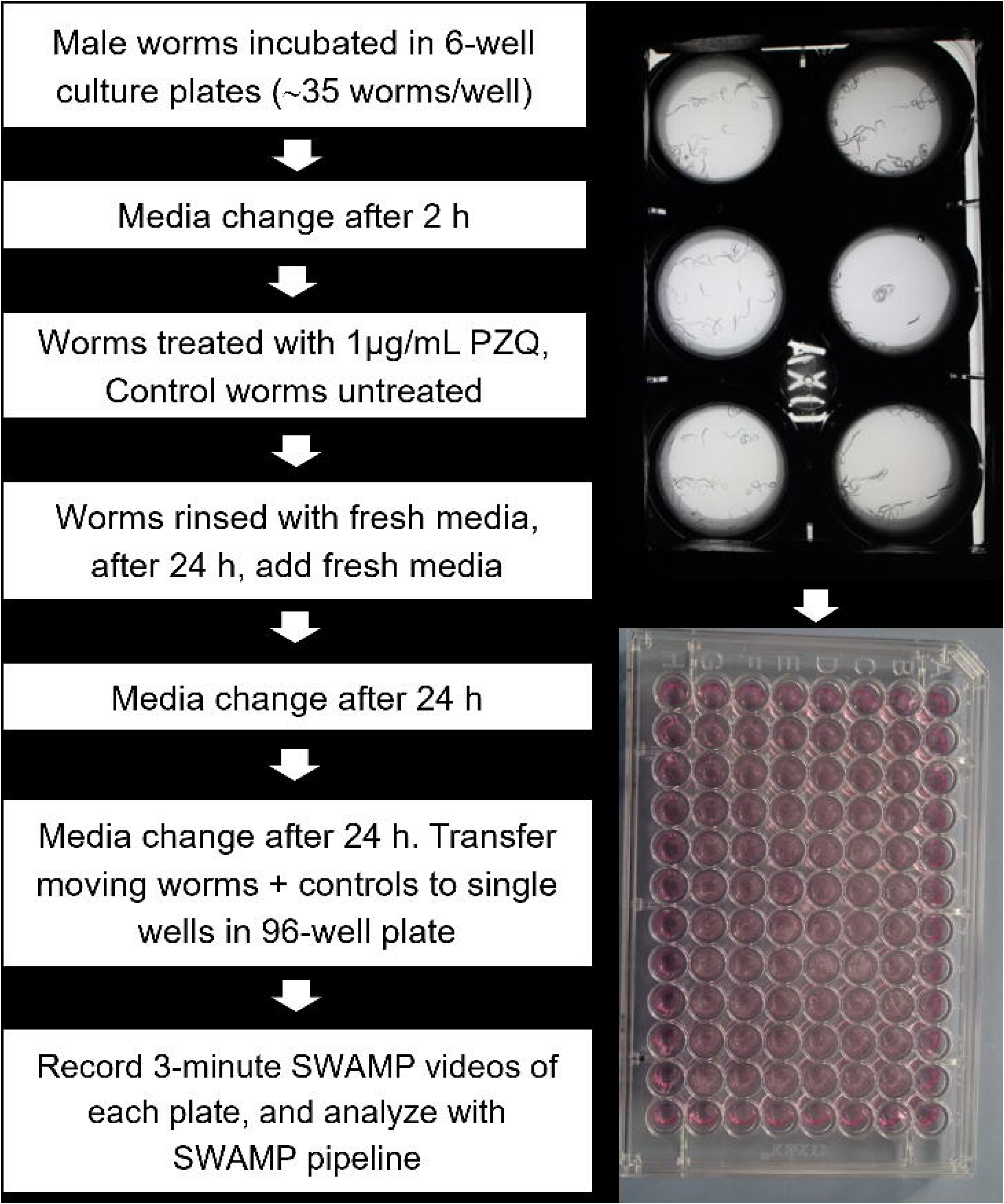
Workflow for conducting single worm movement assays. Field derived adult male worm parasites were cultured in six-well plates, treated with PZQ (or DMSO for the control worms) and after five days of culture, the Single Worm Analysis of Movement Pipeline (SWAMP) box and video camera was used to make a 3-minute video recording of each 96-well plate. This was done in the main Neglected Tropical Disease (NTD) laboratory at the Kenya Medical Research Institute’s Centre for global Health Research, Kenya.

### Infection of snails

We collected fecal samples from a total of 87 *S. mansoni* infected participants (3-34 per village) from 5 hotspot and 1 non-hotspot villages in Siaya, Western Kenya (Table 1). We hatched eggs isolated from pooled fecal samples from each village. We exposed 401-480 *Biomphalaria pfeiferri* snails (2801 total) to *Schistosoma mansoni* miracidia (5 per snail) obtained from the 6 villages. We used laboratory reared snails (n=1893) for most infections; these were supplemented with field collected snails (n=908) for 4/6 villages (Supplementary Table S1). After 7-9 weeks, we screened surviving snails for infection. We found a total of 371 cercariae shedding snails (overall infection rate of 21.4%), representing an estimated 371 parasite genotypes. Infection was variable across the villages, ranging from 110 shedding snails out of 310 surviving (35.5%) for Uhanya village, 84/324 snails (25.9%) for Magawa village, 48/264 snails (18.2%) for Agok village, 31 245 snails (12.7%) for Uyawi village, and 37 445 snails (8.3%) for Dago village. Mumbo, the only non-hotspot included in the study had 61/365 snails (16.7%) infection rate.

**Table 1.**
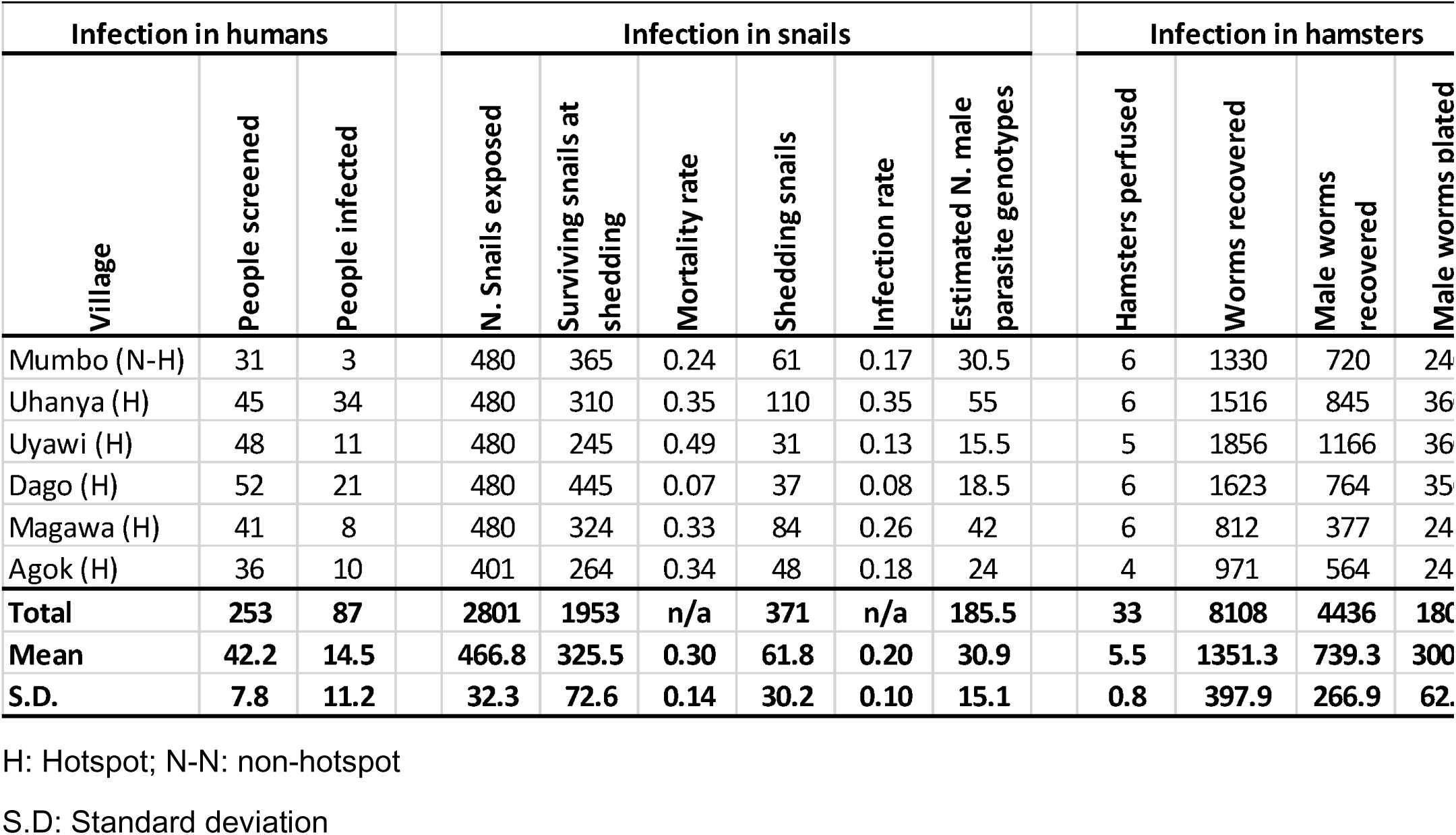
Recovery of adult *S. mansoni* from 6 Kenyan villages. . We hatched eggs from infected people in each village to obtain miracidia, which were then used to infect snails. The snails were lab reared and/or field collected – see supplementary table 1 for details). Note that the number of infected snails provides a conservative estimate of the number of schistosome genotypes and assuming one genotype per snail, while the estimated number of male genotypes is half this number, assuming a 1:1 sex ratio. We collected cercariae from surviving snails (7-8 weeks post-infection), which were mixed in equal proportions to infect hamsters. We perfused adult worms from infected hamsters and used male worms for movement assays to assess evidence for PZQ-resistance.

We experienced considerable challenges achieving high snail infection rates as well as maintaining infected snails due to high mortality rates (0.07-0.49). Laboratory reared snails (maintained for at least one generation in the laboratory) had similar infection rate (GLMM, binomial family, z = -1.174, p = 0.241) and mortality (GLMM, beta-binomial family, z = 0.404, p = 0.686) compared to field-collected snails (Supplementary table S1).

We conservatively assumed that each infected snail harbors a single schistosome genotype. Hence, we had 31-110 schistosome genotypes per village, which equates to 16.5-55 male genotypes assuming a 1:1 sex ratio (Table 1). We are likely to underestimate genotype number because some snails will be infected with >1 parasite genotype. We used the frequency of uninfected snails and binomial assumptions to estimate numbers of snails infected with 1, 2, 3, 4 or 5 genotypes and the mean number of parasite genotypes per snail (Supplementary table 2). The estimated mean number of genotypes per infected snail ranged from 1.03 (Uhanya) to 1.19 (Dago), indicating that our conservative assumption of 1 genotype per snail is reasonable.

### Hamster infection and worm perfusion

For each village, we infected 6 hamsters with pools of 700 cercarial made by combining equal numbers of cercariae from each infected snail. Each of these hamsters was infected with cercarial pools with identical composition. We perfused hamsters after 55 days and separated male and female pools in petri dishes containing DMEM complete media. We recovered 1351 ± 397 (SD) adult worms per village (8108 total), of which 739 ± 267 (SD) were male (4436 total).

### Worm motility assays

<u>Control experiments</u>: We used mixtures of PZQ-ES and PZQ-ER worms to evaluate the efficacy of our movement assay to identify resistant worms (Fig. 3). We identified two moving worms in a pool of 100 worms containing a single PZQ-R worm. The most active worm was shown to be PZQ-ER by PCR-RFLP analysis. We identified 5 moving worms in a pool of 100 worms containing 5 PZQ-ER worms, all of which were PZQ-ER by PCR-RFLP. The movement assay correctly identified all 6 PZQ-R worms (sensitivity 100%, 95% CI 54–100%) and correctly classified 193/194 PZQ-S susceptible worms (specificity 99.5%, 95% CI 97.2–100%). The high specificity suggests this assay is a robust tool for detecting PZQ parasites. However, specificity is <100% so some moving worms detected following PZQ treatment may be false positives.

**Figure 3.**
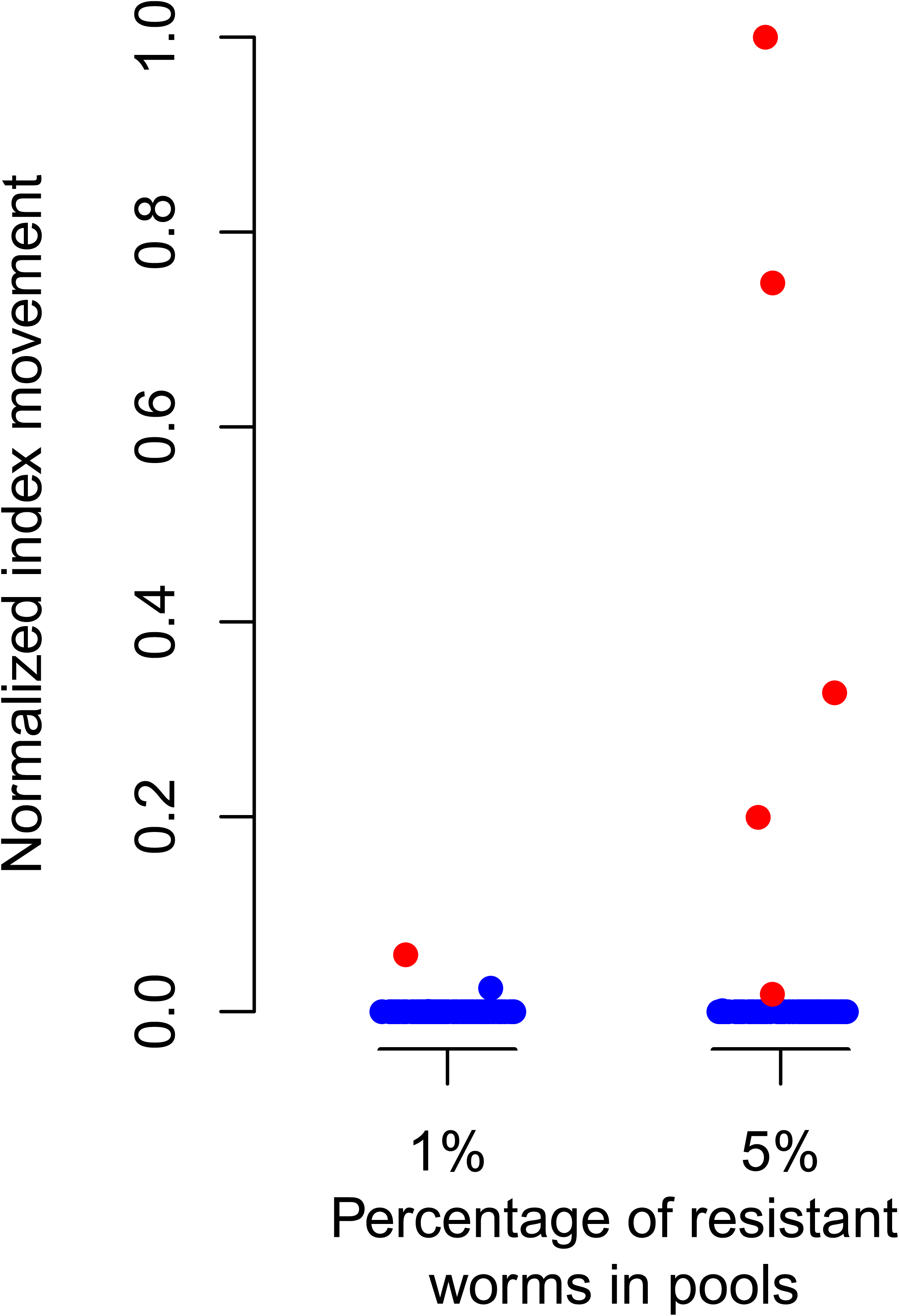
Control experiments to evaluate efficacy of movement assays for detection of PZQ-R parasites. A. we constructed mixtures of 100 worms containing SmLE-PZQ-ER and SmLE-PZQ-ES parasites in the ratio of 1:99 and 5:95. B. Results of movement assays on the SWAMP assay platform. Red circles show PZQ-ER parasites, while blue circles are PZQ-ES parasites. Black circles are PZQ-ES individual for which the genotype was not determined.

<u>PZQ-response in field derived worms</u>: Following hamster infection, euthanasia and perfusion, we screened a total of 1,800 adult male worms, representing an estimated 185 genotypes, for *in vitro* susceptibility to PZQ using the Single Worm Analysis of Movement Pipeline (SWAMP) assay. Overall, we observed 1/1800 moving worms after PZQ-treatment. These worms comprise an estimated 185 genotypes, giving a frequency of recovered worm genotypes of 1/185 (frequency = 0.54%; 95% CI 0.01 - 2.97%, exact binomial). After PZQ treatment, we observed no moving worms among 1,561 worms screened from 5 hotspot villages (Fig. 4). These worms comprise an estimate 155 worm genotypes, hence the frequency of recovered worm genotypes is 0/155 (95% CI 0-2.3%, exact binomial). We observed one weakly moving worm from 240 worms representing an estimated 31 genotypes from the non-hotspot village, giving a frequency of 1/31 (3.2%, 95% CI 0.08-16.6%, exact binomial). The difference between five hotspot and the one non-hotspot village was not statistically significant (Fisher’s exact p = 0.33), although confidence intervals were wide due to low event counts. These results demonstrate just one worm showing weak movement following *in vitro* PZQ exposure. We note that in our control experiments a single PZQ-sensitive parasite was moving following PZQ-exposure, so we cannot rule out that the surviving worm was a false positive, rather than a true PZQ-R parasite.

**Figure 4.**
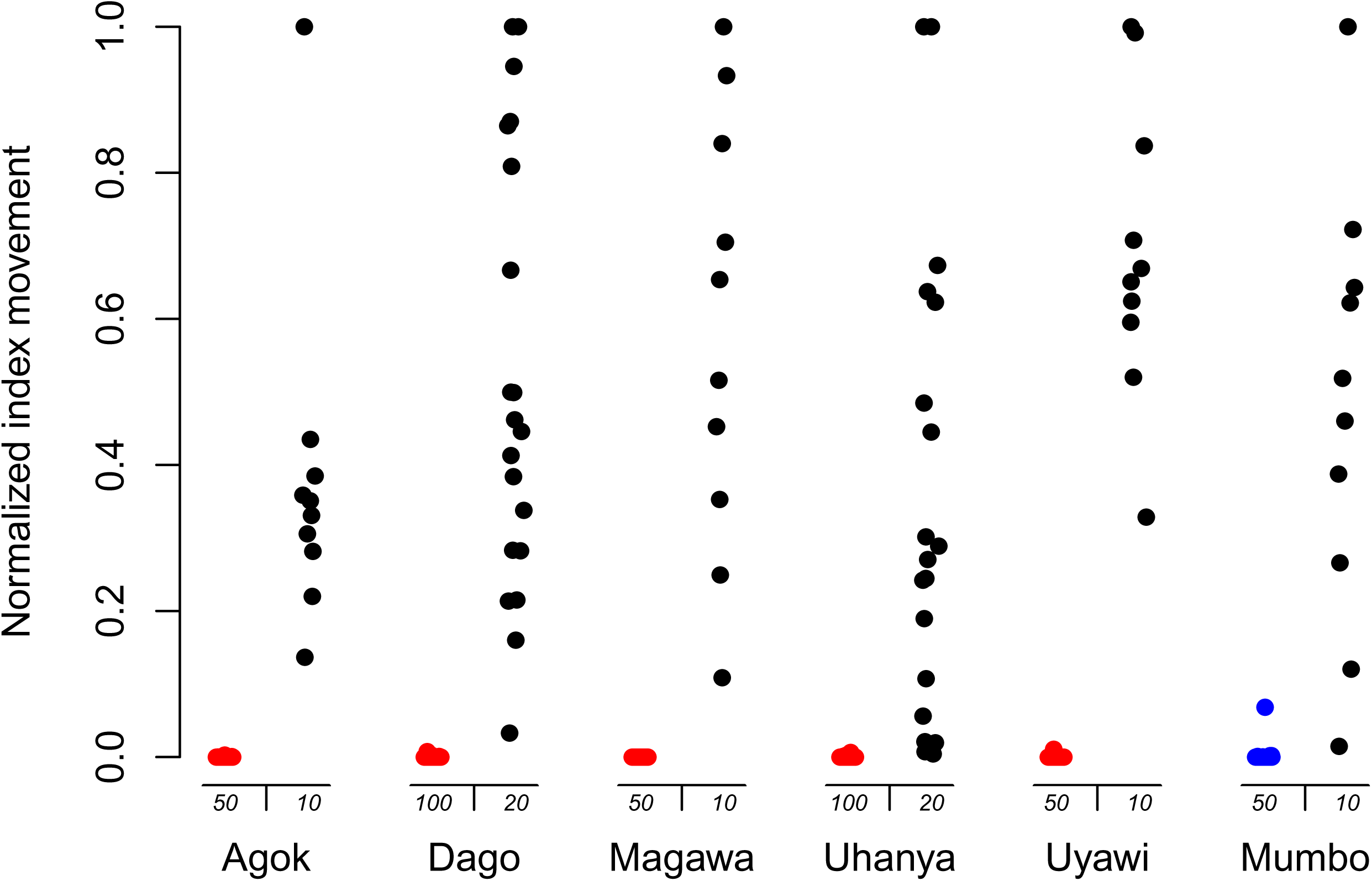
Drug response assays for adult male parasites from 6 Kenyan villages. Normalized movement indexes are plotted, such that the most active worm for each village is set at 1. We observed no movement in PZQ-treated worms from hotspots (red) and one moving worm from the non-hotspot (blue) village. Corresponding controls (black) are actively moving (Kruskal-Wallis, Χ² = 383.52, p-value < 2.2e-16, pairwise Wilcoxon rank sum tests available in supp. Table S3). Numbers under the x-axis indicate number of PZQ-treated and untreated examined for movement.

We included DMSO treated control worms for comparison with PZQ treated worms on each culture plate (see videos deposited in BioImage Archive). The untreated control worms showed significantly greater movement than the PZQ-exposed male worms (Kruskal-Wallis, Χ² = 383.52, p-value < 2.2e-16, pairwise Wilcoxon rank sum tests available in Supplementary Table S3).

## DISCUSSION

### No evidence for PZQ-R in Western Kenya

We screened 1,800 adult male parasites, representing an estimated 185 parasite genotypes and observed a single worm moving post PZQ exposure. Our observation of one resistant worm in an estimated 185 worm genotypes examined has 95% confidence intervals of 0.01 - 2.97%, (binomial confidence intervals). These results suggest that PZQ-R worms are rare or absent in Western Kenyan *S. mansoni* populations, given that we are unable to eliminate rate false positives using the movement assay. We used a PZQ concentration of 1μg/mL for the assays. From our previous work, sensitive laboratory population (SmLE-PZQ-ES) have IC_50_ of 0.19 μg/mL, while resistant laboratory population (SmLE-PZQ-ER) have IC_50_ of 73 μg/mL (Le Clec’h et al., 2021).. Hence, the PZQ concentration is ∼5 times the IC50 value observed for PZQ-sensitive worms in the laboratory. The worms examined originated from 5 hotspot villages and one non-hotspot village, with the single moving worm coming from a non-hotspot village. Our results suggest (i) that PZQ-R is unlikely to explain the persistent high prevalence and intensity observed in hotspot villages in the face of frequent mass treatment with PZQ and (ii) that other features of these locations explain persistent hotspots.

Our phenotypic screen results are further supported by a study of TRPM_PZQ_ sequence variation in miracidia pools collected from 10 villages in Western Kenya, which included the 6 villages studies here. This molecular surveillance study examined allele frequencies of mutations within the 1,695bp TRP box region of TRPM_PZQ_ and found just 3 mutations at > 1% frequency, none of which impacts channel activation by PZQ (Olilah et al, in preparation). Furthermore, allele frequencies of these mutations were not significantly different in hotspot and non-hotspot villages. That we found no putative resistance mutations in the molecular survey is consistent with the conclusions from our phenotypic screen. We caution that these encouraging results from Western Kenya using both phenotypic and molecular surveillance methods should not be automatically extrapolated to other endemic areas. It would be particularly informative to conduct further phenotypic screens for PZQ-R in other locations, particularly in locations where treatment of humans reveals ERR significantly lower than 0.95 (Fukushige et al., 2021).

### Estimation of numbers of parasite genotypes screened

While phenotypic screens allow direct examination of drug response in adult parasites and avoid many of the issues inherent in interpretation of treatment studies in patients, they are laborious to perform. To conduct these assays required establishment and maintenance of both snail and hamster colonies. Furthermore, 3-4 months are required for *S. mansoni* development in *B. pfeifferi* snails (8 weeks) and in hamsters (45 days) to generate adult worms for phenotypic screens. We designed our assays to maximize the numbers of parasite genotypes examined in order to obtain accurate measures of frequencies of resistant parasites within parasite populations from each village. Hence, while we examined 1,800 adult male parasites, these represented an estimated 185 independent genotypes. PZQ-resistance is a genetic trait (Le Clec’h et al., 2021), so genotype number rather than worm number is the appropriate denominator for these phenotypic assays.

We aimed to maximize numbers of adult worm genotypes examined as follows. First, we collected miracidia from multiple patients (mean=14.5; range 3-34) in each village to help ensure good representation of parasites from the population. Prior work on *S. mansoni* population structure from East Africa revealed that that there is limited structure at the level of individual infected patients (Berger et al., 2021; Steinauer et al., 2013). Furthermore, each miracidium larva is genetically unique. Hence, sampling miracidia from several patients minimizes bias of population samples from each village (Steinauer et al., 2013). Second, we infected as many snails as possible (mean=466.8; range 401-480). A severe bottleneck occurs during snail infection. For each village examined, we infected multiple snails with 5 miracidia each: hence the cercariae emerging from each snail will comprise between 1-5 different genotypes. Infection rates were low (8.3-35.5%) suggesting that most snails were infected by just a single miracidium. Assuming infection rates show a Poisson distribution, infected snails should contain on average 1.03-1.19 genotypes in the six villages studied (Supplementary table 2), and numbers of genotypes represented among cercariae emerging from snails may be lower than this. Therefore, our estimate of one parasite clone from each snail provides a conservative estimate.

### Technical challenges with phenotypic screens

Obtaining sufficient numbers of snails for these infections, and the low infection rate (8.3-35.5%) was the major limitation on obtaining large numbers of adult worm genotypes. We used *Biomphalaria pffeifferi* for these assays. We also explored use of two other species, *B. sudanica* and *B. choanomphala*. *B. sudanica* was easier than *B. pffeifferi* to maintain and produce large numbers of snails, but infection prevalence and numbers of cercariae shed were much lower from *B. sudanica* (Lu et al., 2016). Similarly, while *B. choanomphala* also shows higher infection rate than *B. sudanica* (Mutuku et al., 2021), we were unable to rear sufficient numbers of *B choanomphala*, so focused our efforts on *B. pfeifferi*. Development of methods for increasing production and infection rates of snails would simplify future phenotypic screening for PZQ resistance in *S. mansoni*.

One possible approach might be to identify lineages of *B. pfeifferi* with elevated infection rates. *B. pfeifferi* preferentially self-fertilizes (Kengne-Fokam et al., 2016; Mintsa Nguema et al., 2013), so snail populations comprise different lineages of genetically identical multilocus parasite genotypes (MLG). Recent work in Western Kenya suggests that some MLGs show elevated infection rates with trematodes, including *S. mansoni*. (Oduor et al., 2025). We suggest that developing laboratory colonies of an MLG showing high susceptibility to *S. mansoni* could provide an approach to improve infection rates.

### What other factors might explain hotspots?

Our phenotypic survey found a single moving parasite in populations from hotspots and non-hotspots in western Kenya. That suggests that factors other than PZQ-R likely explain the persistent hotspots. These include behavioral, biological, environmental, programmatic, socioeconomic and WASH (water, sanitation, and hygiene) factors (Assaré et al., 2020b, 2020; Kittur et al., 2020; Olilah et al., 2026; Sokouri et al., 2024) Occupational and behavioral factors include duration and frequency of contact with schistosome-infested water, driving transmission. A study in Côte d’Ivoire reported higher frequency and density of human water contact sites in hotspots (Assaré et al., 2020) compared with non-hotspots. Environmental and ecological factors affect the abundance and distribution of the intermediate snail vector, with favorable ecological niches allowing sustained parasite transmission despite ongoing preventive chemotherapy with PZQ (Sokouri et al., 2024). Mutuku et al (Mutuku et al., 2019) found that hotspot villages along the shores of Lake Victoria had higher abundance of the deep-water vector species *B. choanomphala* than non-hotspots, perhaps promoting transmission in these villages. Socioeconomic status determines access to safe WASH infrastructure, and poor access to these has been shown to contribute to persistent transmission (Kittur et al., 2020). Programmatic and intervention-associated limitations are also insufficient to interrupt transmission in hotspots (Kittur et al., 2020). We conclude that factors other than PZQ-R are likely to explain hotspots in Western Kenya.

## MATERIALS AND METHODS

### Study area, study participants and diagnosis

The study was carried out in villages in Rarieda and Bondo sub-counties of Siaya county, western Kenya. This was part of a larger study investigating phenotypic and molecular surveillance of drug resistance in schistosomes in western Kenya carried out between February 2022 and August 2024. Ethical permission for work with humans was from the KEMRI ethical review board (protocol SERU 3218) and Kenyatta University Ethical Review Committee (Protocol no. PKU/2206/I1352). We recruited participants from six villages, five of which had been classified as *Schistosoma mansoni* transmission hotspots by the SCORE study in 2017 (Wiegand et al., 2017), and confirmed in a recent epidemiological survey that hotspots retained significantly higher prevalence (Olilah et al., 2026). A total of 253 participants were initially consented to provide a stool sample for diagnosis of *S. mansoni* infection from the six villages. A single stool sample of about 2 g was provided by each participant. These were then transported in cooler boxes to the Kenya Medical Research Institute’s Centre for Global Heath Research, Neglected Tropical Diseases Laboratory in Kisian, Kisumu for processing. The samples were immediately used to prepare slides by Kato-Katz technique, using two slides per sample. An experienced technician prepared and observed the slides under a microscope for schistosome eggs.

### Miracidia hatching and snail infection

Having identified infected individuals, we collected additional stool samples for purification of eggs and isolation of miracidia. This was done for each of five hotspot and one hotspot village. We isolated parasite eggs by processing fecal samples through 3 layers of sieves (350 μm to 60 μm). Isolated eggs were then placed in petri dishes and exposed to light to allow miracidia hatching.

We established a large breeding colony of *Biomphalaria pfeifferi* snails, which were used as intermediate snail hosts for *S. mansoni*. For infection, we placed snails individually in wells in a 24 well culture plate containing 2 ml of water per well. We exposed each snail to 5 miracidia for ∼5 hours in wells of a 24-well plate to allow sufficient time for infection. The snails were then returned to their aquarium tanks. After 3 weeks they were transferred to opaque tanks to minimize light exposure that would stimulate cercariae shedding and increase snail mortality. We screened for shedding snails in 24 well culture plates 8 weeks after the initial infection, having determined that this was the best time for optimal cercarial shedding. When insufficient snails were available in our laboratory colonies, we collected *B. pffeiferi* snails from field locations. These were screened for shedding snails. We then used uninfected snails for exposure to miracidia as described for laboratory reared snails. Lab and field collected snails were maintained separately to assess mortality and infection rates

We constructed pools of cercariae for infection of hamsters using equal numbers of cercariae from each shedding snail. For example, if 100 shedding snails were available, we added 7 cercariae from each snail to make pools of 700 cercariae to infect each hamster. We made identical cercariae pools for infecting each hamster infected. This procedure was used to maximize the number of parasite genotypes infecting each hamster.

### Hamster infection and perfusion

We established a breeding colony of hamsters at KEMRI (Kisian, Kisumu) using animals procured from breeders or donated from the animal house at KEMRI Nairobi. We used at least 6-week-old hamsters for schistosome infection. Ethical permission for work on hamsters was from the KEMRI ethical review board and Animal care and Use Committee (Protocol 3218) and Kenyatta University Animal Care and Use Committee (PKUA/001/001). On the day of infection, we placed each animal in a bucket containing one inch of warm water for 10-15 minutes to wet their fur to improve infection efficiency. Hamsters were then allowed to wade individually for 2 h in one inch of water containing the cercarial pools (see above) within a closed glass jar with breathing holes in the lid. After 55 days, we perfused the hamsters as previously described (Duvall & DeWitt, 1967), and the collected adult worms were separated into male and female pools in petri dishes containing DMEM complete media (DMEM High glucose – L-glutamine + 15% HiFBS + 10 mL/L antibiotic stock solution (10 UI penicillin / 10 µg/mL streptomycin)).

### Control experiments

We performed control experiments to evaluate the ability of the movement assay to identify individual resistant parasites. To do this, we used SmLE-PZQ-ER and SmLE-PZQ-ES parasite populations generated by laboratory selection. These populations have been purified by marker assisted selection and show more than 377-fold difference in their response to PZQ (Clec’h et al., 2021). We collected adult worms generated during our routine lifecycle maintenance using previously described methods (Jutzeler et al., 2024) under IACUC protocol 1419 MA. We constructed pools of 100 adult male parasites containing SmLE-PZQ-ES and SmLE-PZQ-ER worms in proportions 99:1 and 95:5. The worms were mixed and split in 6 well plates cultures (33 to 34 worms per well). Worms were cultured for 5 days and were all treated with PZQ (see *In vitro* culture and PZQ treatment section). On day 4, we plated all worms in 96-well plates (60 worms per plate). We then quantified their movement using the SWAMP assay to identify worms recovering after PZQ exposure. Videos are available in the BioImage Archive under accession number S-BIAD3516 (DOI: 10.6019/S-BIAD3516).

We selected the 10 worms showing the most movement and genotyped them using a PCR-RFLP assay to determine whether they were SmLE-PZQ-ER or SmLE-PZQ-ES. We extracted single worm DNA using the Chelex method (Criscione et al., 2009). We amplified a 421 bp segment (forward primers: 5’-TCGTAATAAACATGGTCGTC-3’, reverse primer: 5’-TCGACTACAGAATGATGTAA-3’) on chromosome 3 which contains a marker associated to PZQ-response (Clec’h et al., 2021). Each PCR reaction was as follows: 9.325 μL of sterile water, 1.5 μL of 10x buffer, 1.2 μL of dNTP (2.5 mM each), 0.9 μL of MgCl_2_, 0.5 μL of each primer (10 μM), 0.075 μL of Taq polymerase (TaKaRa) and 1 μL of gDNA template. PCR amplification was done using the following program: 95°C for 5 minutes, [95°C for 30s, 55°C for 30s, and 72°C for 1min] × 35 cycles, 72°C for 10 minutes. We then digested PCR amplicons with Mse I enzyme. Reaction was as follows: 6.5 μL of sterile water, 1 μL of 10x CutSmart buffer (NEB), 0.5 μL enzyme (5U/µL; NEB), 2 μL of PCR amplicon. Incubation was as follows: 37°C for 1h, 65°C for 20 minutes. The digested products were visualized by electrophoresis on a 2% agarose gel.

### *In vitro* culture and PZQ treatment

We placed live male worms from the perfused hamsters into sterile 6-well culture plates. Each well contained ≤ 35 worms in 3.5ml of DMEM complete media. Plates were maintained in an incubator at 37°C with 5% CO_2_ and media was warmed up to 37°C before use. On day 1, we renewed the media in each well after 2 hours of incubation and incubated the plates for 24 hours under the same conditions. On day 2, we prepared the treatment media (DMEM complete media containing 1 μg/mL of PZQ sampled from a stock solution of PZQ in DMSO at 10 mg/mL) and the control media (DMEM complete media containing the same volume of DMSO as the drug stock volume sampled for the treatment media). We changed the media with the treatment media in 5 wells, while we changed the media with the control media in one well. On day 3, we removed media and used fresh media to rinse the worms three times to remove residual PZQ. We then added fresh media and incubated the plate for a further 24 hours. On day 4, the media was changed with fresh media and the plate replaced in the incubator. On day 5, 48 hours after drug treatment, we inspected each well under a binocular microscope. We transferred moving worms from the PZQ-treated wells to a 96-well plate (1 worm per well) containing 100 µL of fresh media in each well. We transferred non-moving worms based on visual inspection from the PZQ-treated wells as negative controls to reach a total number of 50 worms. Moving and non-moving worms were randomly distributed on the 96-well plate. We also added DMSO-treated worms in the first row of each 96-well plate (B2-B11) as positive controls. In total, up to 60 worms were plated, leaving the border wells filled with media only to mitigate evaporation. The plate was incubated for one hour. Each plate was then placed in the Single Worm Analysis Pipeline (SWAMP) box and a 3 minute and 20 second video recording made. The design of the SWAMP box is available on Zenodo (DOI: 10.5281/zenodo.21300824). We analyzed the videos using the SWAMP pipeline (<u>github.com/fdchevalier/SWAMP</u>) to quantify worm movement. Briefly, the pipeline samples video frames every 5s, aligns them, isolates wells and compares pixel change to estimate worm movement. We compared movement between PZQ-treated worms and DMSO-treated worms (positive control). Videos are available in the BioImage Archive under accession number S-BIAD3520 (DOI: 10.6019/S-BIAD3520) (Hartley et al., 2022).

### Number of genotype estimation

We calculated the number of parasite genotypes examined for PZQ resistance as follows: Each snail was infected with 5 miracidia. Therefore, shedding snails will produce cercariae of 1-5 genotypes. We conservatively assumed that each snail was shedding a single genotype – this will underestimate the true numbers of genotypes. Hence, if 100 snails were shedding, we estimated that cercarial pools (constructed with equal numbers of cercariae from each shedding snail) would contain 100 parasite genotypes. We assumed an equal parasite sex ratio, so a cercarial pool of 100 genotypes was estimated to contain 50 male genotypes.

### Statistical analysis

Infection rates and survival were compared between lab-reared and field snails using Generalized Linear Mixed Models (GLMM). For both response variables, infection rate and survival were modelled as binomial proportions in R, with group (lab-reared vs. field) as a fixed effect and location as a random effect to account for variation across experimental sites. Models were fitted using maximum likelihood estimation (Laplace Approximation) via the lme4 package (Bates et al., 2015). Model fit was assessed by calculating the overdispersion ratio as the ratio of residual deviance to residual degrees of freedom. Residual diagnostics were further evaluated using simulated residuals from the DHARMa package, including tests for uniformity, overdispersion, and zero-inflation. Where overdispersion was detected, a beta-binomial model was fitted using the glmmTMB package (McGillycuddy et al., 2025).

We used the frequency of uninfected snails and binomial assumptions to estimate numbers of snails infected with 1, 2, 3, 4 or 5 genotypes, and the mean number of parasites per infected snail. This assumes that snails are equally susceptible to infection, and the number of cercarial genotypes shed from a snail is equal to the number of miracidia that successfully infected that snail.

We calculated the sensitivity and specificity of our movement assay to detect PZQ-R parasites in control experiments using a confusion matrix. We computed 95% confidence interval using binomial distribution.

We calculated frequencies of worm genotypes that were still motile after PZQ exposure and corresponding 95% confidence intervals. We tested differences in worm movement with a Kruskal-Wallis test, followed by pairwise Wilcoxon rank sum tests with Bonferroni correction to identify group differences.

We performed all statistical tests in R (version v4.1.2: (*R: The R Project for Statistical Computing*, n.d.)). We considered differences to be statistically significant when p-values < 0.05.

The code used for all analyses is available on Github (https://github.com/fdchevalier/PZQ_response_Kenyan_worms).

## DECLARATIONS

### Ethics approval and consent to participate

The study protocol was reviewed and approved by both the Kenyatta University Ethical Review Committee (Protocol no. PKU/2206/I1352) and the Kenya Medical Research Institute’s Scientific and Ethical Review Unit (SERU), protocol number 3218. All adult participants gave informed consent to take part in the study, while minors gave assent with their guardians signing informed consent on their behalf.

### Authors’ contributions

EMN, FDC, WL,TJCA designed the experiments. JO and EMN consented participants, collected and processed samples, infected and shed snails, infected and perfused hamsters and carried *in vitro* assays and recorded SWAMP videos. PO performed hamster perfusions and helped with *in vitro* cultures. MM conducted the control experiment, which included hamster infection and perfusion, worm cultures and treatment, SWAMP videos recording, and molecular genotyping. FDC and WL supervised the control experiment and analyzed SWAMP videos. TJCA and EMN wrote the first version of the manuscript. All authors edited the manuscript and approved the final version.

## Acknowledgements

We thank the study participants and community health workers from the 6 villages for their involvement with this project. We thank Kathrin Jutzeler, Stephanie Nordmeyer, and Neal Platt for snail maintenance. We thank Alison Whigham, Oscar Rodriguez, and the vivarium technicians for their assistance with hamster husbandry.

